# Diversification dynamics of the Palaeozoic actinopterygian radiation

**DOI:** 10.1101/2025.04.14.648759

**Authors:** Joseph Flannery-Sutherland, Struan Henderson, Sophie Fasey, Sam Giles

**Affiliations:** University of Birmingham

**Keywords:** fossil sampling bias, spatial standardisation, birth-death process, speciation, extinction, origination

## Abstract

Ray-finned fish are the most speciose vertebrate group today, but the dynamics of their early diversification are contentious. Their fossil record suggests a first radiation in the Carboniferous following the Late Devonian mass extinction events. Conversely, recent phylogenetic hypotheses imply a radiation originating in the Late Devonian, but lack the taxonomic breadth required to robustly test this. This necessitates phylogeny-free inference of actinopterygian diversification rates from fossil occurrences, itself challenging due to complex systematics, incomplete occurrence databases, and severe spatiotemporal sampling biases. Here, we analyse a comprehensive dataset of Palaeozoic actinopterygian genera and species using approaches that accommodate spatial and temporal sampling variation. We detect elevated actinopterygian diversification in the Late Devonian, with substantially greater lineage survival across the Hangenberg extinction event than indicated by the raw fossil record. Surprisingly, we detect no positive shifts in origination across the event, refuting previous hypotheses of explosive actinopterygian radiation in its wake. Instead, cryptic survival of diversified lineages appears responsible for the robust signal of increased diversity across different geographic scales in the Carboniferous. Nonetheless, these trends are overwhelmingly driven by low palaeolatitude Euramerican fossil assemblages, highlighting the ongoing spatial limitations of the actinopterygian fossil record.

## 1. Introduction

The early diversification of ray-finned fish (Actinopterygii) presents an intriguing evolutionary puzzle. Following their divergence from lobe-finned fish (Sarcopterygii) in the Silurian [1] and in sharp contrast to their sister clade, actinopterygians persisted at low diversity and ecological prominence for nearly 50 million years, represented by a handful of undisputed occurrences through the Early and Middle Devonian, alongside fragmentary taxa known only from scales with contested taxonomic affinities [2,3]. The Late Devonian was punctuated by the Kellwasser crises and Hangenberg event, mass extinction pulses marked by reef collapse and marine anoxia, across which vertebrate faunas previously dominated by placoderms and sarcopterygians were rapidly supplanted by a more contemporary fauna composed of chondrichthyans and taxonomically and morphologically diversified actinopterygians [4-10].

Previous studies have hypothesised that the Hangenberg event was directly responsible for the sudden inflection in the macroevolutionary trajectory of actinopterygians [4-6,11,12], casting their Carboniferous explosion of diversity as a classic model of adaptive radiation following mass extinction. This interpretation of their empirical fossil record appears borne out by phylogenetic hypotheses of fossil taxa, which infer a paucity of lineages to have survived the event and bring the origin of crown group actinopterygians in line with the Devonian-Carboniferous boundary [11,13]. A recent report of a Devonian actinopterygian nested within an otherwise Carboniferous clade [13] raises an alternative hypothesis, however: that cryptic actinopterygian diversification began in the Devonian under markedly different ecological conditions and with much higher lineage survivorship into the Carboniferous than previously thought. [13,14].

The poor quality of the early actinopterygian fossil record presents a substantial challenge to evaluating whether their early Carboniferous diversification was fuelled by explosive radiation of a few lineages following the Hangenberg event, or by wider survival of lineages which diverged during the Devonian [13]. Many taxa are morphologically conservative, known from only fragmentary specimens, or both, hindering confident anatomical characterisation and inclusion in morphological data matrices. In turn, phylogenetic analyses of taxonomically depauperate, morphologically incomplete matrices have typically recovered extensive polytomies between early actinopterygian taxa and uncertain relationships between Palaeozoic lineages and crown actinopterygians more generally, including candidates for Devonian lineages seeding diversification in the Carboniferous [12,16-21]. Fossil occurrence databases have also failed to keep up with the complex and extensive systematics of Palaeozoic actinopterygians where lengthy history of study, largely unaided by modern imaging techniques and often of character-poor specimens, has resulted in a web of problematic taxonomic delineations and revisions [10]. Consequently, no analysis of early actinopterygian fossil diversity from either occurrence-based or phylogenetic approaches has utilised an up-to-date compilation of all described taxa.

Recent efforts have tackled the lack of high-quality fossil occurrence databases but have also highlighted the severe sampling biases plaguing the early actinopterygian fossil record through geological time and across geographic space [5,9,10,16]. As such, it is unknown whether their sharp increase in apparent diversity in the early Carboniferous might be driven by spatiotemporal sampling biases in their fossil record, or if their increased geographic extent genuinely reflects their evolutionary radiation at this time. Patterns of relative diversity are themselves useful indicators of macroevolutionary success but are generally uninformative of the rates of origination and extinction responsible for changing diversity. A range of abiotic or biotic mechanisms can drive rate variation [22], with implications for the processes which may have shaped early actinopterygian diversity in the ecological context of the Late Devonian versus the early Carboniferous. Competitive exclusion by other gnathostomes for example could have suppressed origination, while environmental vulnerability might have resulted in high extinction, but these rates have not yet been quantified in Palaeozoic actinopterygians.

Two broad strategies have been proposed to overcome spatial sampling biases. The first uses diversity estimates from local subsamples of the fossil record where spatial sampling variation is minimised between subsamples [23-26], allowing informative average trends to be drawn from these local estimates. The second uses larger, regional datasets which are spatially standardised through time to capture taxon stratigraphic durations rather than just pointwise samples of diversity [27-29]. Origination and extinction rates can then be robustly estimated from these datasets using birth-death-sampling models which accommodate the phylogenetic linkage of a set of taxa subsampled from clade’s total diversity, enabling rate estimation that accommodates the unsampled ghost lineages required to tackle the problem of Late Devonian diversification, but without the need for a formal phylogenetic tree [30-33]. In this paper, we investigate patterns of diversity and diversification rates in early actinopterygians using an expanded fossil occurrence database coupled with these complementary analytical strategies to elucidate their rise to prominence through the Palaeozoic. We characterise their sampling and diversity through the Palaeozoic to understand the extent to which their fossil record is biologically informative versus driven by sampling bias, particularly the potential role of geographic sampling variation through time. We then focus on whether exceptional diversification dynamics were responsible for their early Carboniferous radiation or whether their radiation was instead rooted in the Late Devonian.

## 2. Materials and methods

### 2.1 Fossil occurrence data

We assembled an occurrence dataset of all valid early actinopterygian genera and species described up to February 2024 within the Paleobiology Database (PBDB). This comprises 2142 Lochkovian (Early Devonian) to Changhsingian (Late Permian) genus level actinopterygian occurrences, of which 1644 (76.7%) are classified to species level, across 907 stage- to substage-constrained PBDB collections with a mean collection age uncertainty of 4.1 Ma (median 2.8 Ma, median absolute deviation 2.1 Ma). We also included 212 Early Triassic occurrences across 62 PBDB collections to minimise edge effects in our study window. Generally, these collections are specific geological sites from which fossils have been collected, although a smaller number represent regional-scale faunal assemblages or estimated sampling locations for certain taxa. Rather than using arbitrary equal length intervals or formal chronostratigraphic divisions in our downstream diversity analyses, we used the formation binning approach of Dean et al. [34] to optimise the resolution of our downstream diversity analyses without exceeding the practical limits imposed by occurrence stratigraphic uncertainty, thus avoiding a false degree of temporal precision in our results. Full details on the assembly, cleaning and binning of our dataset are available in the supplementary information.

### 2.2 Spatial subsampling

We used the minimum spanning tree (MST) length of the occurrence data in each time bin to quantify the spatial sampling extent of the actinopterygian fossil record through the Palaeozoic, as MST length captures both the geographic coverage and spread of a dataset and closely correlates with several alternative spatial metrics [23]. As there is substantial variation in MST length through time (see Results, Fig. 1B), we applied two spatial subsampling schemes to the temporally binned occurrence data to reduce bias from the species-area effect prior to estimation of diversity and diversification rates, and to deconvolve global actinopterygian diversity trends into their local geographic components.

**Figure 1.**
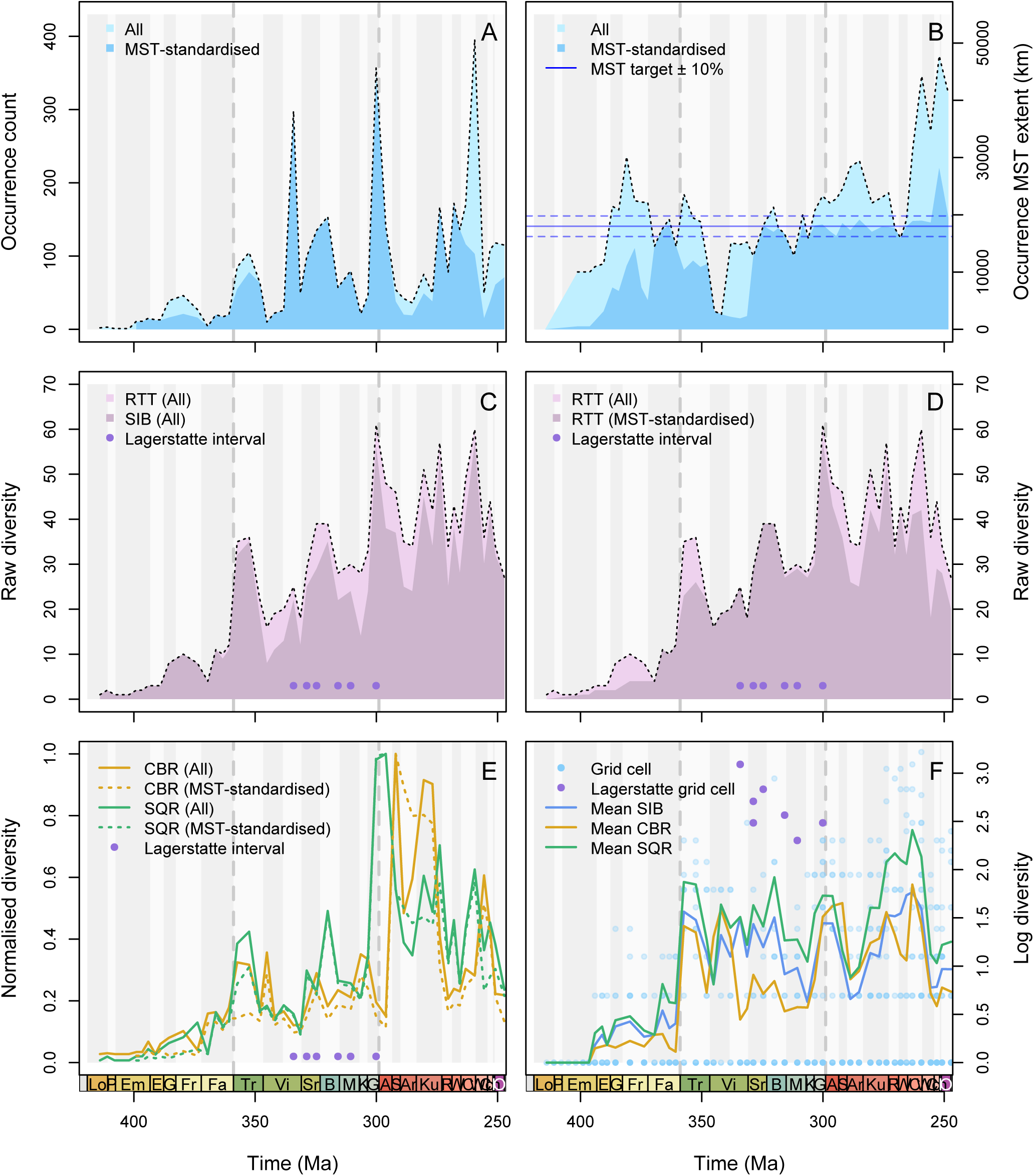
Sampling and diversity in Palaeozoic actinopterygian fossil occurrence record. The coloured ribbon and grey bars demarcate geological stages, with division of the Viséan into informal early and late substages. Vertical dashed lines indicate end of period. Stage abbreviations from left to right as follows: Lo = Lochkovian, P = Pragian, Em = Emsian, E = Eifelian, G = Givetian, Fr = Frasnian, Fa = Famennian, Tr = Tournaisian, Vi = Viséan, Sr = Serpukhovian, B = Bashkirian, M = Moscovian, K = Kasimovian, G = Gzhelian, A = Asselian, S = Sakmarian, Ar = Artinskian, Ku = Kungurian, R = Roadian, W = Wordian, C = Capitanian, Wc = Wuchiapingian, Ch = Changhsingian, I = Induan, O = Olenekian. **A.** Number of fossil occurrences through time. **B.** Geographic extent of fossil occurrences measured as the length of their minimum spanning tree (MST). **C.** Global actinopterygian genus richness measured from the total dataset using sampled-in-bin (SIB) and range-through-time (RTT) metrics. **D.** RTT global actinopterygian genus richness before and after standardisation by MST length. **E.** Sampling standardised global actinopterygian diversity estimated by coverage-based rarefaction (CBR) and Squares (SQR), before and after standardisation by MST length. **F.** Local actinopterygian genus richness measured in equal area grid cells using SIB, CBR and SQR metrics, plus mean grid cell richness through time.

Our first scheme reduces fluctuations in the global extent of an occurrence dataset through time by subsampling its MST to a target length (18000 km) between time bins, using the method of Flannery-Sutherland et al. [27]. This provides the taxon stratigraphic durations required for estimating diversification rates as well as diversity, whilst preserving their phylogenetic diversification signal by minimising taxonomic exclusion [27]. Choice of target MST length presents a trade-off between standardisation efficacy versus data exclusion between time bins [27]. As more stringent thresholds would have resulted in unacceptably high exclusion, spatial extent of the available data in some intervals naturally falls below our target length, either because actinopterygians were geographically restricted at that time in their evolutionary history or due to poor sampling. The former reason may be plausible for the earliest portion of their fossil record, but the latter is likely responsible for later deficient intervals, so we avoid drawing any biological conclusions from these even though we still estimate diversity and diversification rates across them. Our second scheme does not provide data suitable for rate estimation but avoids the limitations of the MST approach by aggregating temporally binned collections within equal area hexagonal grid cells generated by the *icosa* R package [35] to provide samples of local diversity, followed by calculation of mean grid cell diversity through time.

### 2.3 Diversity estimation

We estimated bin-wise genus richness (= diversity herein) from our total and spatially subsampled datasets using sampled-in-bin diversity (SIB), coverage-based rarefaction (CBR) and extrapolation by Squares (SQR), in addition to range-through-time diversity (RTT) for our total and MST-subsampled datasets. CBR and SQS are more suited to estimating true diversity patterns and have previously been applied to Palaeozoic actinopterygians [8] but not within spatially controlled frameworks that mitigate geographically biased sampling. SIB makes no corrections for uneven sampling between time bins and RTT only ensures that the minimum bound on diversity is recovered, but these metrics still provide useful indications of sampling quality in the fossil record, and permit comparison with counts made by previous studies. We favoured genus-level analysis as genera are considered more robust to variation in sampling and taxonomic practice, reducing the potential for highly uneven abundance distributions which can undermine the reliability of CBR [36], although it is not a true substitute for species level diversity [37]. SIB and RTT were calculated using the `divDyn()` function of the *divDyn* R package [38]. CBR was implemented using the `estimateD()` function of the *iNEXT* R package [39] with a sampling quorum of 0.5, and SQR calculated directly in R using the equation of Alroy [40].

### 2.4 Birth-death-sampling models

We estimated Palaeozoic actinopterygian origination and extinction rates to investigate the timing and magnitude of any rate shifts, using the total occurrence dataset which represents their known diversity, as well as our MST-standardised dataset to additionally control for the confounding effects of spatial sampling variation. Traditional methods for inferring diversification rates from fossil occurrence data (e.g., second-for-third or three-timer metrics [41,42]) are often unreliable when confronted by clades with preservation rates and high turnover [43]. Instead, methods which treat fossil occurrence data as the product of a birth-death-sampling process are more robust [44]. These methods are also insensitive to uneven taxon abundance distributions that impact CBR or SQR, so we infer origination and extinction rates at genus and species levels, although we note that the latter level still displays problematic over-splitting of morphologically conservative wastebasket genera (e.g., *Rhadinichthys*, *Elonichthys*, *Amblypterus*, *Paramblypterus*).

We employed two models with complementary sets of assumptions: a hierarchical birth-death-sampling model [30-32] and the skyline fossilised birth-death range model [33,44] hereafter referred to by their respective implementations in the software packages PyRate [31] and RevBayes [45]. Briefly, the key model differences are, respectively, in statistically-derived versus user-prescribed timings of shifts in origination and extinction rates, and the assumption of complete versus incomplete sampling of all fossil lineages that ever existed. As such, PyRate avoids imposing any rate shifts across the Hangenberg event *a priori*, while RevBayes relaxes the unrealistic assumption of complete lineage sampling. Both PyRate and RevBayes ultimately produce estimates of origination, extinction, and sampling rates through time, permitting direct comparison between their different model assumptions. Full details of the implementations and parameters for all rate analyses are available in the supplementary information.

## 3. Results

### 3.1 Palaeozoic actinopterygian sampling trends

The first putative actinopterygian fossils occur in the Lochkovian (Fig. 1A). Bin-wise counts of actinopterygian occurrences are low through the Devonian until a two-to-threefold increase in the Tournaisian (∼ 355 Ma, Fig. 1A). Prominent spikes occur late in the Viséan (∼340 Ma), through the Bashkirian and Gzhelian (∼320 and 302 Ma), and in the Wuchiapingian (∼258 Ma), producing a saw-toothed sampling pattern through the Palaeozoic (Fig. 1A). The spatial extent of their fossil occurrences also varies through time, falling below our target spatial extent of 18000 km in the Early and Middle Devonian, and in the Viséan (Fig. 1B). Subsampling by MST length resulted in relatively minor loss of occurrences from the dataset but was generally effective in reducing fluctuations in spatial extent from the Serpukhovian (∼330 Ma) with extent in most bins falling within ±10% of our target (raw MST length standard deviation = 8942 km; subsampled MST length standard deviation = 6801 km; Fig. 1B), aside for a slight deficit early in the Bashkirian, and in the Changhsingian (∼253 Ma) where spatial extent slightly exceeds the target (Fig. 1B).

Spikes in occurrence sampling correlate with total SIB diversity (Spearman’s rank correlation coefficient = 0.90, p < 0.0001), while the relationship with their total spatial extent is weaker, but still significant (Spearman’s rank correlation coefficient = 0.34, p < 0.05), with Devonian extents comparable to those throughout the Carboniferous and Permian despite substantially more counts in the latter. Controlling for bias under the species-area effect by MST subsampling reduces actinopterygian SIB diversity in the Frasnian (∼370 Ma), in the Tournaisian and through the Permian, but SIB diversity through the rest of Palaeozoic is generally unaffected. RTT diversity reduces sawtooth trends in SIB diversity only slightly (Fig. 1C), with SIB showing deficits relative to RTT in the Viséan, Bashkirian, Moscovian (∼310 Ma) and Artiniskian (∼285 Ma).

### 3.2 Palaeozoic actinopterygian diversity trends

Global trends in RTT diversity closely track with estimates from CBR (Spearman correlation coefficient = 0.82, p < 0.0001) and SQR (Spearman correlation coefficient = 0.97, p < 0.0001), with little change between the total and MST-standardised datasets (Figs. 1D, 1E). Diversity accrued slowly through the Devonian but rose by around an order of magnitude across the Devonian-Carboniferous boundary. Some diversity loss occurred into the early Viséan before a second peak at the beginning of the Pennsylvanian, decline through the remainder of the Carboniferous, then a general trend of rising diversity through the Early Permian. This trend terminated with a drop in diversity in the Middle Permian, followed by a final spike in the Changhsingian. The major exceptions to this correspondence between raw and sampling-corrected diversity trends, however, are the patterns of relative richness across the Permian-Carboniferous boundary, where SQR diversity peaked sharply in the Gzhelian while CBR diversity peaked in the Asselian (∼298 Ma; Fig. 1E)

At local geographic scales, there is a regime shift from a low to a high mean grid cell diversity across the Devonian-Carboniferous boundary recorded by SIB, CBR and SQR metrics (Fig. 1F). SIB and SQR are highly similar throughout the study interval, with small fluctuations through the Tournaisian to the Bashkirian, a slight decline through the Moscovian and Gzhelian, recovery across the Carboniferous-Permian boundary, a second decline through the Artinskian, progressive rise through the remainder of the Early to Middle Permian, then a final decline through the Late Permian (Fig. 1F). CBR diversity instead declined in the late Viséan until the recovery across the Carboniferous-Permian boundary, while the progressive diversity rise through the Middle Permian recovered by SIB and CBR was punctuated by decline through the Roadian and Wordian. The prominent Gzhelian and Asselian spikes in global diversity recovered by SQR and CBR respectively are more muted but remain apparent at the local scale (Fig. 1F).

### 3.3 Palaeozoic actinopterygian diversification rates

Actinopterygian genus and species origination (λ*_g_*, λ*_s_*) and extinction (μ*_g_*, μ*_s_*) rates are sensitive to spatial sampling extent and birth-death-sampling model assumptions (Figs. 2, 3). In our PyRate results, μ*_g_* was virtually constant from the Devonian to the Capitanian stage of the Middle Permian. In the total dataset, μ*_g_* increased twofold at the end of the Capitanian then rose through the remainder of the study window, compared to a sixfold increase at the end of the Changhsingian across the Permian-Triassic boundary in the spatially standardised dataset. In the complete dataset, λ*_g_*was constant through the Devonian, sharply decreased in the early Tournaisian into a period of constancy punctuated by a weak, transient increase at the end of the Carboniferous, then increased threefold in Lopingian (Fig. 2D). After spatial standardisation, the sharp decrease in λ*_g_* instead took place across the Devonian-Carboniferous boundary (Fig. 2D). The net result was a piecewise downshift in positive mean diversification rate across the Devonian-Carboniferous boundary, transient increases at the end of the Carboniferous and the Early Permian, then an increasingly negative rate through the Lopingian (Fig. 2F). Species-level rates from PyRate are more volatile than at the genus level (Fig. 3), with diversification rate increasing very gradually through the Devonian, then fluctuating between positive and negative regimes from the Carboniferous onwards (Fig. 3F).

**Figure 2.**
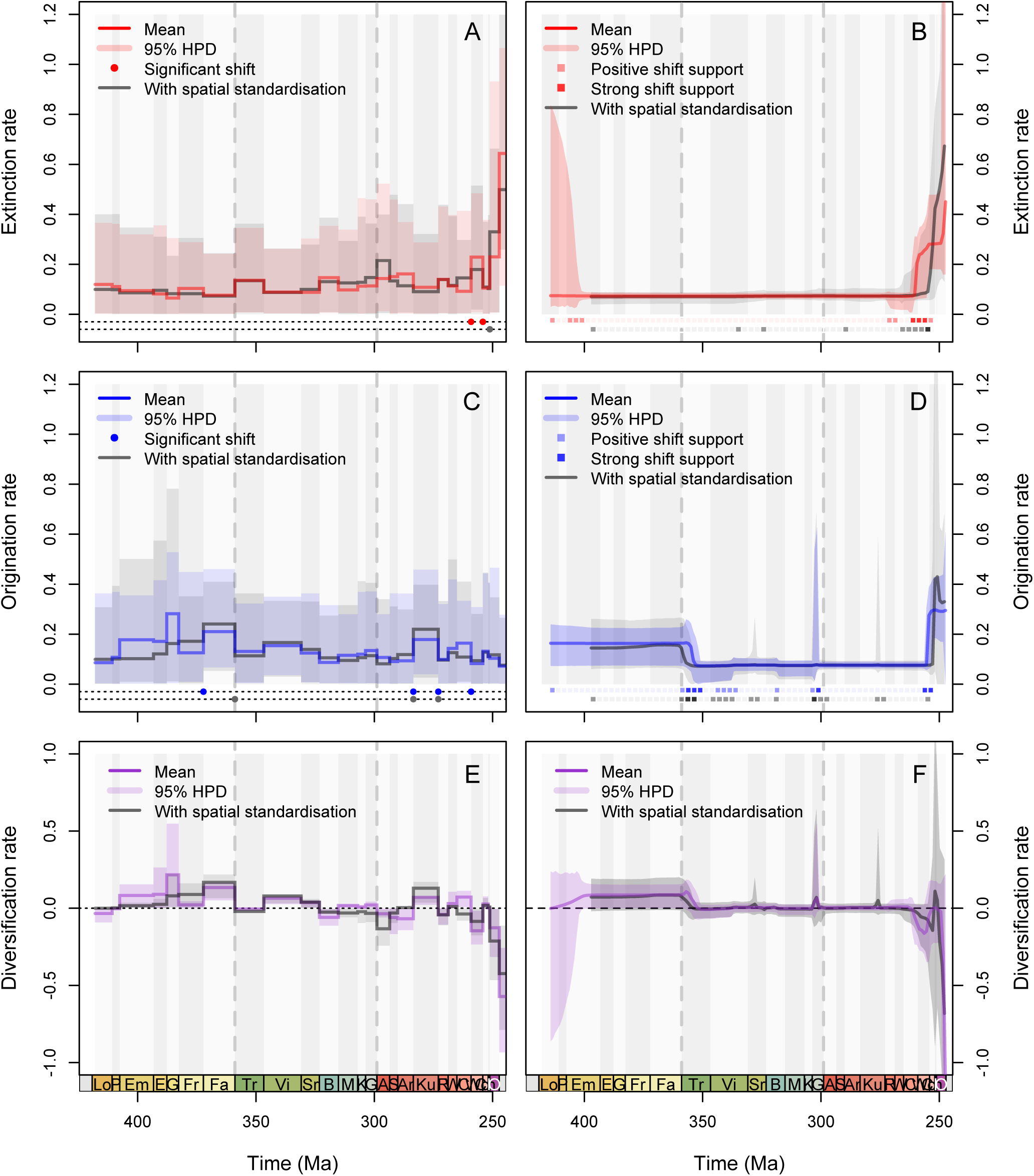
Genus-level diversification dynamics of Palaeozoic actinopterygians. Origination (λ*_g_*) and extinction (μ*_g_*) rates, resultant net diversification rates, and 95% highest posterior densities were estimated under two different birth-death-sampling processes using Bayesian inference. Results in colour are from the total dataset, while results in grey are from a subsample of the dataset following spatial standardisation by minimum spanning tree length. Statistical support for the significance of extinction and origination rate shifts is displayed at the bottom of each plot. The coloured ribbon and grey bars demarcate geological stages, with division of the Viséan into informal early and late substages. Vertical dashed lines indicate end of period. Stage abbreviations as in Fig. 1. **A.** RevBayes μ*_g_*. **B.** PyRate μ*_g_*. **C.** RevBayes λ*_g_*. **D.** PyRate λ*_g_*. **E.** Genus diversification rate from RevBayes. **F.** Genus diversification rate from PyRate.

**Figure 3.**
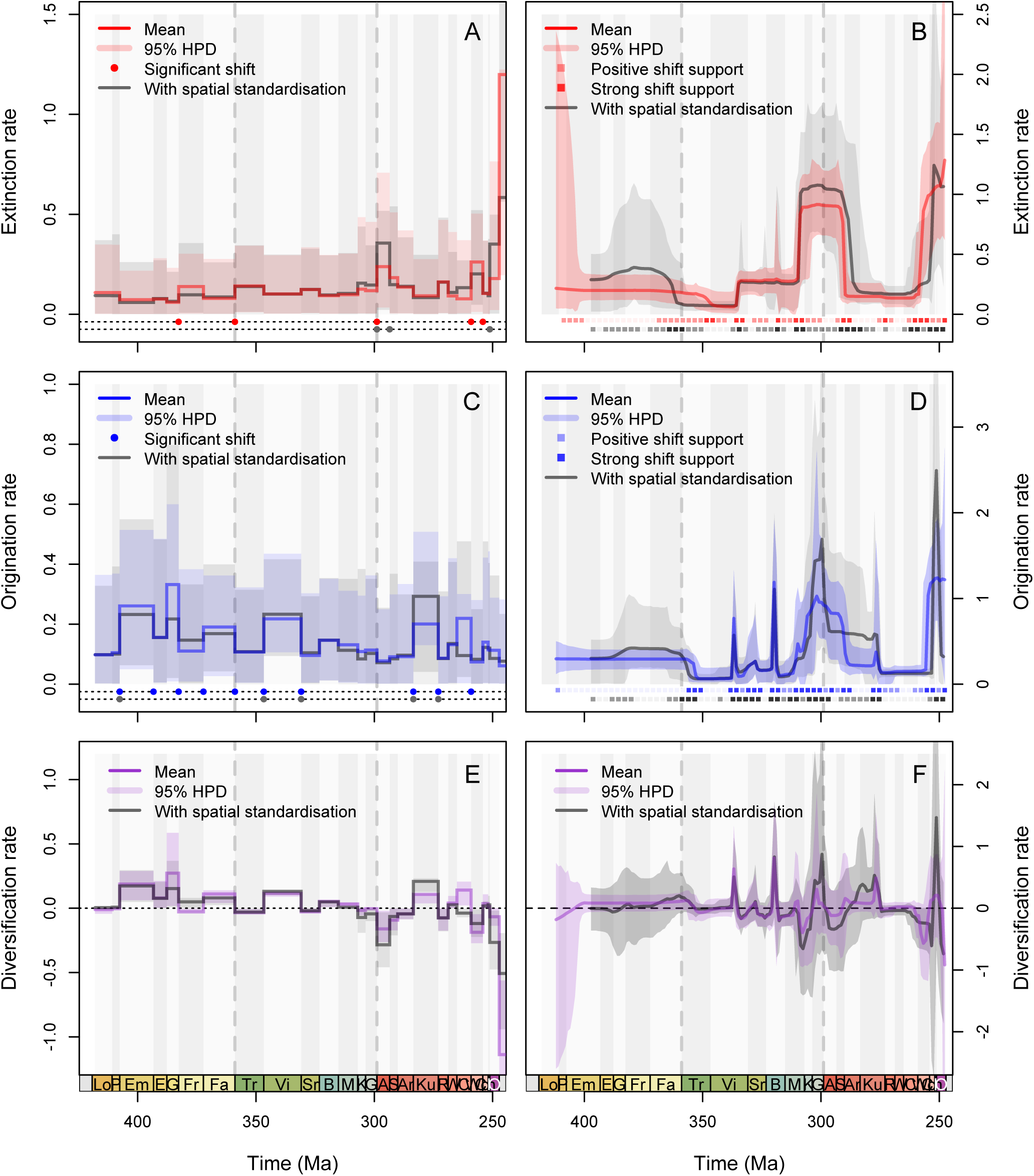
Species-level diversification dynamics of Palaeozoic actinopterygians. Origination (λ*_s_*) and extinction (μ*_s_*) rates, resultant net diversification rates, and 95% highest posterior densities were estimated under two different birth-death-sampling processes using Bayesian inference. Results in colour are from the total dataset, while results in grey are from a subsample of the dataset following spatial standardisation by minimum spanning tree length. Statistical support for the significance of extinction and origination rate shifts is displayed at the bottom of each plot. The coloured ribbon and grey bars demarcate geological stages, with division of the Viséan into informal early and late substages. Vertical dashed lines indicate end of period. Stage abbreviations as in Fig. 1. **A.** RevBayes μ*_s_*. **B.** PyRate μ*_s_*. **C.** RevBayes λ*_s_*. **D.** PyRate λ*_s_*. **E.** Species diversification rate from RevBayes. **F.** Species diversification rate from PyRate.

PyRate found no evidence for a statistically significant increase in diversification rate at the Devonian-Carboniferous boundary at either the genus or species levels before or after spatial standardisation (Figs. 2B–2D, 3B–3D). Instead, diversification rates generally showed a modest decline at the onset of the Carboniferous (Figs. 2E-2F, 3E–3F), driven by a statistically significant decrease in origination rate, an elevated rate in the late Pennsylvanian driven by elevated origination, and a sharp decline towards the end of the Permian driven by a statistically significant increase in extinction rate. Statistically significant rate shifts in origination and extinction are more prevalent at the species level compared to the genus level and are also more prevalent after spatial standardisation, but many of these are positively rather than strongly supported and of negligible magnitude. Exceptions are spikes in the 95% highest posterior density in λ*_s_* in the Serpukhovian and the end of the Kungurian, although the change in mean rate is again minimal (Fig. 3D).

Origination and extinction rates from RevBayes are more dynamic than those from PyRate and display greater uncertainty (Figs. 2, 3), with estimated sampling rates also differing substantially to those of PyRate and suggesting a high prevalence of incomplete lineage sampling (Fig. S1). μ*_g_* declined gently through the Early and Middle Devonian, fluctuated through the Late Devonian, rose then fell in the Tournaisian and Viséan respectively, rose and fell again at the beginning of the Pennsylvannian epoch of the Carboniferous, then continued to rise for the remainder of the Carboniferous and into the Early Permian (Fig. 2A). μ*_g_* dropped again at the end of the Late Permian, spiked in the Wuchiapingian, then spiked even more strongly across the Permian-Triassic boundary. These trends remain after spatial standardisation, apart from a more rapid rise in μ*_g_* through the Pennsylvanian, then a more sudden decline in the Early Permian (Fig. 2A). In contrast to extinction, λ*_g_*showed stepwise increases through the Early and Middle Devonian, a drop in the Frasnian stage of the Late Devonian, a spike in the Fammenian stage at the end of the Devonian, then another decline in the Tournaisian (Fig. 2C). λ*_g_* rose slightly during the Viséan, then decreased to Early Devonian levels through the remainder of the Carboniferous and into the Early Permian (Fig. 2C). Spikes in origination punctuated the Kungurian and Wordian stages of the Early and Middle Permian respectively, but rates remained generally low otherwise (Fig. 2C). Trends are again largely unchanged following spatial standardisation, aside for a continued stepwise rate increase through the Devonian. These rates resulted in intervals of strongly positive diversification in the Middle Devonian, Fammenian, Viséan and Kungurian, diversification close to zero in the Tournaisian, and further fluctuations between positive and negative regimes through the remainder of the Palaeozoic until a crash at the end of the Permian (Fig. 2E). Species-level dynamics from RevBayes show similar trends to those at genus level with more prominent spikes in origination and extinction rates, although the absolute magnitudes of species-level rates are generally lower than those recovered by PyRate (Fig. 3), and the timing of rate increases and decreases for μ*_s_* in the Pennsylvanian are concordant before and after spatial standardisation, with a peak in extinction rates centred in the Asselian (Fig. 3A).

Bayes Factor tests marked very few RevBayes rate shifts as statistically significant. At the genus level, the only significant extinction rate shifts took place at the end of the Permian, but the timings of individual shifts are discordant before and after spatial standardisation (Fig. 2A), while significant shifts were also detected after spatial standardisation around the Permian-Carboniferous boundary at the species level. Following spatial standardisation, significant decreases in λ*_g_* took place at the end of the Devonian, and at the ends of the Early and Middle Permian (Fig. 2C), while a significant increase took place at the beginning of the Kungurian. As with PyRate, species level rate shifts are more prevalent. Shifts in extinction rate following spatial standardisation were detected at the beginning and end of the Permian (Fig. 3A), while shifts in origination rate are dispersed through the Devonian, Mississippian and Kungurian (Fig. 3C).

## 4. Discussion

### 4.1 Signals of sampling versus macroecology

The positive correlation between geographic extent and observed diversity in the Palaeozoic actinopterygian fossil record indicates that spatial sampling bias affects their diversity trends. Despite this, raw actinopterygian diversity patterns through the Devonian and Carboniferous were largely unchanged by spatial standardisation procedures (Fig. 1D). At face value this might suggest that their fossil record during this interval is well sampled enough to adequately record their diversity patterns, despite fluctuation in sampling intensity and spatial extent, but a more realistic interpretation is that trends are so heavily biased towards their intensely sampled low latitude Euramerican fossil record [8,9] that spatially standardised diversity recovers the trend of their total fossil record. Our expanded occurrence database, incorporating all early actinopterygian taxa as of our cut-off date remains subject to this fundamental limitation in the spatial sampling structure of their fossil record. Consequently, it is not strictly possible to identify global trends in actinopterygian diversity, necessitating the potentially flawed assumption that the diversity patterns from our predominantly Euramerican dataset are representative of their global dynamics.

Despite bearing most of their Palaeozoic fossil record, the oldest actinopterygian occurrences come from outside of Euramerica and are themselves geographically disparate (the South China Block and Brazilian Paraná Basin [46-47]), even though actinopterygians must necessarily have originated in a discrete location. Similarly, these occurrences are from marine facies, but the oldest undisputed Euramerican actinopterygian, *Cheirolepis trailli* from the Middle Devonian, is from lacustrine deposits of the Orcadian Basin of Scotland. Actinopterygians clearly dispersed widely early in their evolutionary history, including from the marine to the freshwater realm, yet there is essentially no fossil record of these transitions [17]. Alongside their conspicuous absence in the Silurian in contrast to their sarcopterygian sister group, these geographic and environmental disparities belie the sampling deficits plaguing their early fossil record. Indeterminate actinopterygian material offers tantalising glimpses into this conspicuous biogeographic gap [9], but its incompleteness prevents adequate characterisation of their wider diversity during the Devonian and continues to highlight the anatomical and morphological challenges facing phylogenetic and occurrence-based approaches to their diversification dynamics. Even so, the abundance of well-sampled Silurian and Devonian sites which record the proliferation of placoderms and sarcopterygians in both marine and freshwater ecosystems suggests that actinopterygians were genuinely rare during this interval in their evolutionary history, partly explaining the paucity of their early fossil record [9]

While spatial extent, and total and sampling-corrected diversity in the actinopterygian fossil record increased from the Devonian to the Carboniferous, this pattern is borne out by average diversity across equal area grid cells, suggesting that this proliferation was a genuine biological trend, rather than an artefact of spatiotemporal sampling biases. There is clearer evidence for undersampling during the early Viséan and Moscovian, where declines in spatial sampling extent correspond to declines in diversity even after MST-subsampling, besides greater deficits between SIB and RTT metrics (Fig. 1C). Sharp crashes in diversity or diversification rates during these intervals are not supported, however, by diversity trends from our hexagonally binned subsamples or results from PyRate or RevBayes (Figs. 1F, 2A–2C, 3E–3F). We therefore predict that future sampling within known or undiscovered early Viséan localities should reveal substantially greater actinopterygian diversity than is currently observed.

Actinopterygian diversity patterns can be partly isolated from variation in spatial sampling extent by focusing on their consistently sampled Euramerican data, but this record may still be prone to other biases, particularly lagerstätten effects. This does not appear to impact diversity patterns observed across from the Famennian to the Tournaisian. Multiple sites bearing exceptional actinopterygian specimens preserved in three dimensions or which have yielded substantial quantities of compressed, but complete actinopterygian specimens within well-preserved community assemblages more generally, however, correspond high diversity intervals through the late Viséan (Glencartholm, Granton), Serpukhovian (Manse Burn, Bear Gulch), Bashkirian (Pennine Coal Measures), Moscovian (Mazon Creek) and Gzhelian (Montceau Les Mines). Lagerstätte intervals may display inflated diversity resulting from disproportionately complete preservation of past assemblages, with exceptional preservation additionally increasing potential for sampling of taxonomically and phylogenetically diagnostic characters [48,49]. Consequently, Euramerican lagerstätten may still bias diversity estimates even after accounting for variation in sampling effort, although lagerstätten effects on diversity are spatially and taxonomically variable [50]. Conceivably, the role of lagerstatten in provisioning exceptionally preserved, character-rich specimens [51] may also impact phylogenetic estimation of early actinopterygian diversification even in intervals lacking noticeable peaks in diversity (e.g., the Kinney Brick Quarry in the Kasimovian).

Permian diversity patterns are more challenging to interpret due to conflicting signals between metrics and spatial scales. Sampling extent is higher compared to the Carboniferous, corroborating previous findings [8,9] and correlating with generally high diversity throughout the period, although diversity is largely undiminished when sampling extent is controlled for (Fig. 1D-1E). High diversity in the Early Permian is also recovered by sampling-corrected metrics before and after spatial standardisation, but the global estimates are abnormally high compared to the rest of the dataset (Fig. 1D–1E), with local diversity estimates showing a more modest diversity rise across the Carboniferous-Permian boundary. A potential explanation is extensive historical collection of small-bodied, morphologically conservative actinopterygian specimens in lacustrine deposits spanning the Asselian and Sakmarian stages of the Early Permian across Europe [52-54], from which many taxonomically suspect genera and species have been described. In turn, highly skewed global abundance distributions during this interval may affect the performance of CBR and SQR, as local diversity patterns do not display such extreme spikes in diversity (Fig. 1F).

At face value, Early Permian reductions in predominantly low latitude local diversity lends circumstantial support to the hypothesis that loss of equatorial coastal and freshwater habitat areas during the assembly and aridification of Pangaea may have curtailed rising actinopterygian diversity [8] or at least in its well-sampled Euramerican component. Diversity reductions did not continue through the remainder of the Permian, however, suggesting that actinopterygians remained locally successful components of marine and freshwater ecosystems. Nonetheless, potential confounding effects from wastebasket taxa or over-splitting of specimens from historic European and Siberian collections cannot be excluded in this instance, emphasising the importance of ongoing and future taxonomic revision to order to ascertain reliable patterns of Early Permian actinopterygian diversity [55,56].

### 4.2 Actinopterygian diversification through mass extinction

The Late Devonian was affected by the two-phase Kellwasser crisis at the end of the Frasnian and the Hangenberg event at the end of the Famennian. The Kellwasser crisis had a more significant impact on reef ecosystems [57], of which actinopterygians were a part (e.g., the Frasnian Gogo Formation of Western Australia [20]). Uncorrected, unstandardised diversity decreased in the early Famennian compared to the Frasnian, but this decrease is lost when sampling corrected metrics or spatial standardisation is employed, potentially due to exclusion of actinopterygians from the Gogo Formation during MST subsampling. Apparent impacts across other vertebrates have been ascribed to back-smearing of lineage-wise extinctions due to poor sampling in the Famennian [4]. While occurrence counts for actinopterygians are indeed higher in the Frasnian, PyRate and RevBayes correct this effect and we find no evidence for a significant increase in extinction rates at either the species or genus levels through the Frasnian (Figs. 2, 3). This corroborates previous inferences that the stratigraphically brief Upper and Lower Kellwasser events had only minor impacts on vertebrate faunas [4]. Resolving the precise effects of these ∼0.1 Ma-duration events on actinopterygians, however, will require section level analysis to achieve a suitable degree of stratigraphic precision.

Previous authors have posited that an explosive radiation of actinopterygians took place in the early Carboniferous [4-6, 11,12]. We find, however, that while actinopterygian diversity increased during this interval across a range of spatial scales, there is no statistical evidence for a significant increase in diversification rate after the Hangenberg event. PyRate returns only weak evidence for increased diversification in the Famennian (Fig. 2E, 3E), but their early Carboniferous rise in diversity remains apparent (Fig. 1C–1F). This suggests that there must have been sufficient lineages present in the Devonian combined with high survivorship into the Carboniferous to facilitate rapid growth in diversity without any clear shifts in diversification rate. PyRate indicates that the true stratigraphic ranges of at least 33 known actinopterygian taxa may have spanned the Devonian-Carboniferous boundary (Fig. 4), highlighting the potential extent of their cryptic diversification and survivorship.

**Figure 4.**
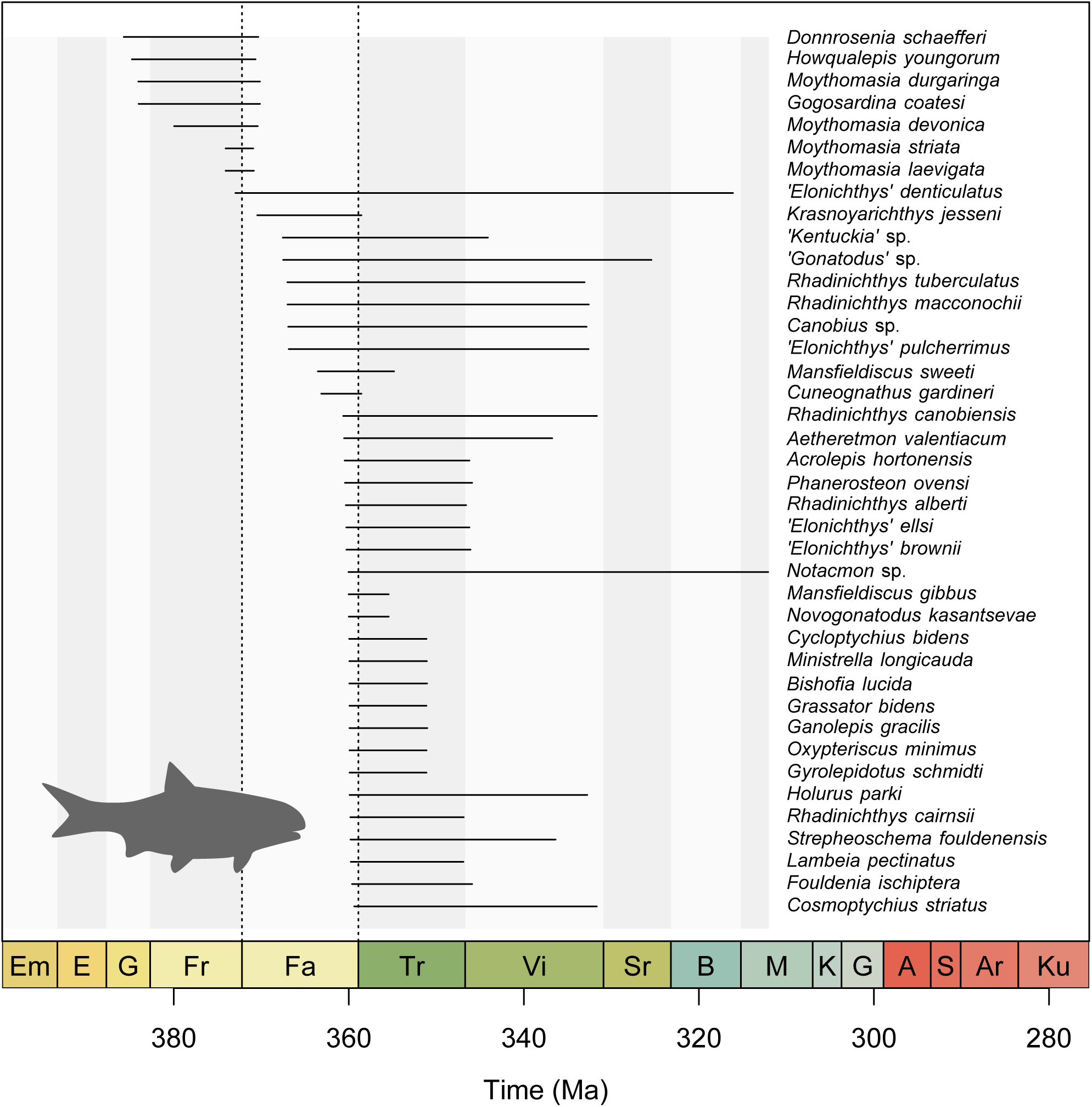
Stratigraphic ranges of actinopterygian taxa spanning the Hangenberg event. Sampling-corrected stratigraphic ranges were estimated using PyRate. The vertical dotted lines mark the timing of the Kellwasser and Hangenberg events at the Frasnian-Famennian and Devonian-Carboniferous boundaries respectively. The coloured ribbon and grey bars demarcate geological stages. The reader is referred to the first figure caption for stage abbreviations. Quote marks indicate suspected polyphyletic wastebasket genera. Silhouette of *Acrolepis* by Tree of Life App.

In some cases, apparent survivorship could be artefactual, arising from backwards extension of wastebasket taxa which share conservative fusiform morphologies. For example, *Kentuckia* and *Gonatodus* are both recovered as boundary crossers, but comparison between their respective Devonian and Carboniferous species strongly suggests that they are polyphyletic [17,58]. In other cases, however, boundary crossers display substantial morphological variation and are also geographically taxonomically disparate [4,59], suggesting that their survivorship and rapid morphological innovation represent the wider diversification dynamics that underpinned early actinopterygian success. Some early Carboniferous taxa retain Devonian-grade anatomy, showing that survivorship was not strictly coupled with morphological innovation [14]. High morphological diversity and abundance displayed by actinopterygians in Tournaisian marine and freshwater deposits [4,60,61], along with evidence for Carboniferous-grade anatomies in some Late Devonian actinopterygians [13,62], indicates that the rapid independent acquisitions of novel body forms across multiple lineages may span a more protracted phase of environmental perturbation and turnover in vertebrate communities that commenced prior to the Hangenberg event [63].

While RevBayes does not estimate true lineage durations, it can relax the assumption of complete species sampling during inference of origination and extinction rates, with lower sampling rates compared to PyRate supporting the expectation that their observed fossil diversity heavily under-samples their true diversity (Fig. S1). Under this relaxed assumption, RevBayes supports a statistically significant increase in genus and species origination rates in the Famennian before and after spatial standardisation, providing the elevated diversification rates needed to generate the cryptic diversity in our proposed evolutionary scenario. Devonian actinopterygian occurrence and taxon counts remain low and so their scarcity through most of this interval appears genuine. There is a conspicuous drop in occurrence counts in the late Famennian following spatial standardisation, however, and so the impression of low diversity prior to the Hangenberg event may partly reflect poor sampling. Indeterminate fragments of Late Devonian and early Carboniferous actinopterygians are also suggestive of their unappreciated taxonomic diversity during this critical interval in their early history [60,64,65]. Complex taxonomic histories and imprecise assignment of numerous actinopterygian specimens to wastebasket taxa likely masks a greater pool of Palaeozoic species diversity than is currently appreciated, while overzealous splitting of anatomically identical specimens will conversely overinflate diversity estimates (e.g., [66]). Future description of actinopterygian taxa in the Famennian or improved sampling of early Carboniferous taxa in phylogenetic trees should address this deficit, particularly given the relative understudy of small bodied, poorly preserved Late Devonian specimens, some of which display Carboniferous-grade morphologies [62]. Cryptic diversification and poor sampling during the Famennian are not mutually exclusive explanations for the apparent disconnect between elevated diversification in the Late Devonian and elevated diversity in the Tournaisian, as a response lag between an increase in rate and its resultant product may be further exacerbated when the latter is also under-sampled.

Results from PyRate and RevBayes both indicate cryptic diversification during the Late Devonian, with respective decreases and increase in origination and extinction rates in the early Carboniferous. This corroborates recent phylogenetically-derived inferences that the recognition of diverse, morphologically distinct actinopterygian lineages in the Carboniferous is predicated on anatomical divergences and lineage separations rooted in the Late Devonian [13] even if Devonian actinopterygian disparity was superficially constrained [67]. These patterns suggest that competitive exclusion by other gnathostome clades may not have been a prevalent mechanism shaping early actinopterygian diversity as, if this were the case, suppression of origination rates would be expected. Preserved gut contents indicate that Devonian actinopterygians preyed upon a taxonomically diverse range of small bodied, mobile prey [67]. This ecological generalism may help explain their stable extinction rates through the Kellwasser and Hangenberg crises by insulating them from turnover and collapse of other vertebrate groups, with previous work showing their similar apparent insensitivity to ecological perturbation during other mass extinction events [68]. Carboniferous decline in origination rates may instead reflect growing intraclade competition as actinopterygians increased in abundance and ecomorphological diversity. Increased extinction rates in the same interval may be a passive component of their greater diversity as the number of lineage terminations will still rise even if proportional extinction rates remain constant. Nonetheless, it could also reflect increased extinction risk in actinopterygian lineages with specialist morphologies and lifestyles that resulted in a naturally greater degree of environmental sensitivity, even though the early Carboniferous was a time of relative stability.

## 5. Conclusions

Birth-death modelling does not support exceptional shifts in actinopterygian diversification rates in the early Carboniferous, but sampling-corrected diversity patterns robustly support a marked increase in the number of actinopterygian taxa during this interval. A possible solution to this apparent contradiction is cryptic diversification of actinopterygians in the Late Devonian coupled with high lineage survivorship across the Hangenberg event. If this scenario is accurate, then formal phylogenetic analyses incorporating a greater proportion of the total known diversity of Palaeozoic actinopterygians and improved sampling of their Famennian fossil record should recover these early divergences. Future analytical gain will also come from resolution of Carboniferous wastebasket taxa through fundamental anatomical redescription. Concurrent critical revision and rejection of dubious scale-based taxa will help to focus efforts on portions of each wastebasket where resolution is even feasible, while at least some wastebasket taxa comprise specimens amenable to CT scanning [58], including three-dimensional specimens of *Kentuckia*, *Gonatodus* and *Rhadinichthy* where taxonomic resolution through robust, detailed anatomical characterisation is possible.

Regime shifts in actinopterygian diversity at local and continental scales observed in the early Carboniferous nonetheless mark the Hangenberg event as a turning point in their macroevolutionary history, with Carboniferous actinopterygians showing wider geographic distributions and ecomorphological diversity compared to in the Devonian [2]. Their increase in taxonomic diversity may not have been driven by exceptional diversification rates, but their change in ecological prominence was still profound. A possible explanation for this is high survivorship of morphologically conservative, cryptic Devonian taxa and ecological release during the Hangenberg event resulting in the decoupling of taxonomic and ecological diversity, a pattern which is seen elsewhere in the fossil record across mass extinction boundaries [69]. Disparity analysis supports morphological conservatism amongst Devonian actinopterygians [67], but at least some the survivors may have displayed higher morphological diversity than is currently appreciated. Further work is therefore needed to ascertain how ecological release versus intraclade competition may have shaped their subsequent evolutionary radiation.

## Acknowledgements

XXX, XXX. and XXX. were funded by a Royal Society Dorothy Hodgkin Research Fellowship (XXX) to XXX. XXX was funded by a Royal Society Enhancement Award to XXX. (grant no. XXX). We thank Emma Dunne, Richard Butler and Matt Friedman for helpful discussions, and three anonymous reviewers whose comments greatly improved the quality of this manuscript.

## Author contributions

XXX and XXX conceived the study. XXX compiled the actinopterygian database at the core of this work which was expanded by XXX. XXX and XXX uploaded the occurrences to the PBDB. XXX carried out all analyses and wrote the manuscript with input from all authors.

## Data availability

All fossil occurrence data used in this work is publicly available through the PBDB. All code scripts and data used to run our analyses are available in the paper electronic supplement (10.6084/m9.figshare.28789472). PyRate and RevBayes are freely available online (github.com/dsilvestro/PyRate; revbayes.github.io/download).

---

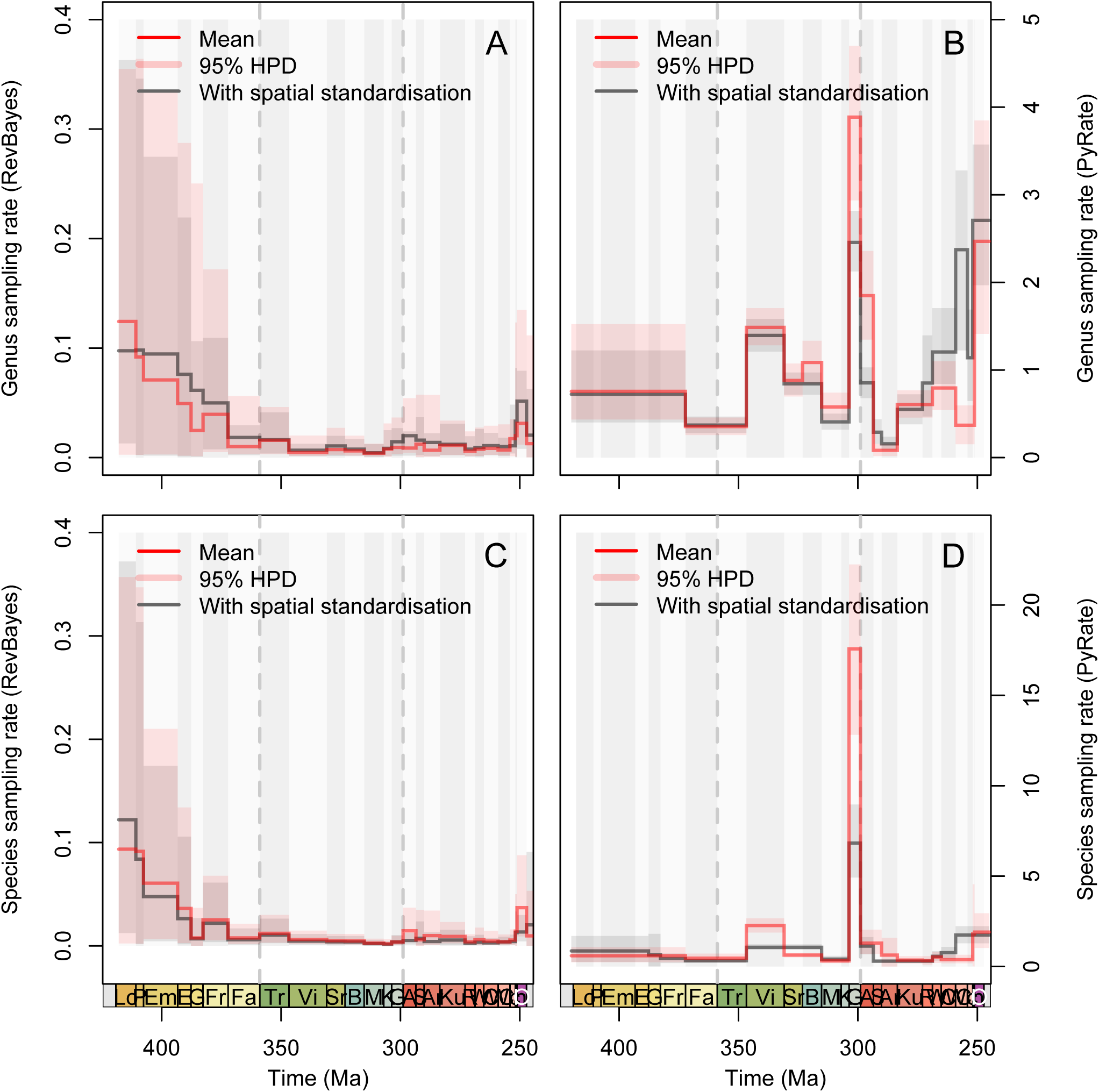

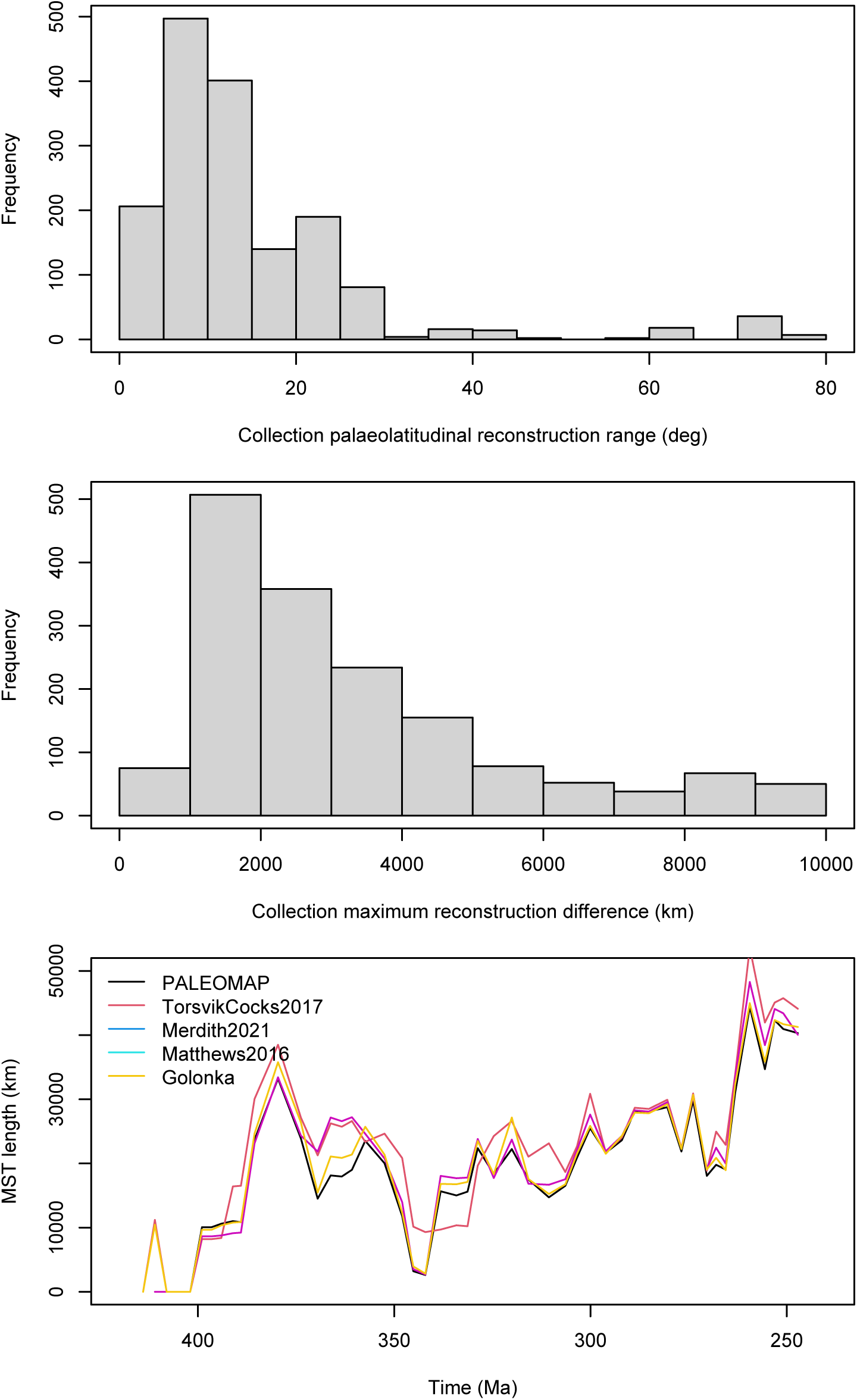

